# Deep Volumetric Localization Super-Resolution Microscopy

**DOI:** 10.1101/2025.05.08.652845

**Authors:** Keyi Han, Xuanwen Hua, Tianrui Qi, Zijun Gao, Xiaopeng Wang, Shu Jia

## Abstract

Super-resolution microscopy, particularly localization-based methods, necessitates careful balancing of optical complexity, computational demands, and user accessibility. Conventional strategies typically adopt either deterministic or learning-based approaches, overlooking opportunities to leverage their synergistic strengths. In this work, we introduce deep volumetric localization microscopy (VLM), a super-resolution methodology that integrates instrumental and algorithmic advancements for high-fidelity 3D single-molecule imaging. VLM employs a wavefront-optimized light-field configuration to capture single-molecule data, while a cascaded neural network reconstructs 3D volumes and extracts molecular coordinates at a 10 nm and 25 nm localization precision in the lateral and axial dimensions, respectively, across an imaging depth exceeding 5 µm. Unlike existing methods, VLM is trained exclusively with system-aware intrinsic point-spread functions, bypassing dependencies on external imaging modalities or sample-specific data training. We validate VLM across diverse biological specimens, demonstrating hardware simplicity, data efficiency, and minimal phototoxicity. We anticipate VLM will overcome current limitations in fluorescence microscopy, empowering broader advancements in biomedical research.

## INTRODUCTION

Over the past two decades, super-resolution microscopy (SRM) techniques have overcome the physical diffraction barriers inherent to traditional optical microscopes, unveiling sub-diffraction details with unprecedented clarity^1-3^. These advancements, spanning dimensions and scales, have enabled the dissection of molecular and cellular mechanisms underlying both physiological and pathological states, thereby offering critical insights for basic and translational biological discoveries^4,5^.

Contemporary SRM methodologies leverage diverse physical and computational principles to surpass the diffraction limit^6,7^. For instance, stimulated emission depletion microscopy techniques (e.g., STED^8^, RESOLFT^9^, MINFLUX^10^) manipulate molecular emission states to achieve resolution beyond conventional limits^2^. Structured illumination microscopy (e.g., SIM^11^, ISM^12^) employs patterned illumination to capture high spatial frequencies, which are subsequently recovered through computational reconstruction^13^. Single-molecule localization methods (e.g., STORM^14^, (f)PALM^15,16^, PAINT^17^) exploit the stochastic activation and precise localization of individual fluorophores^18^. Expansion microscopy (e.g., ExM^19^) physically enlarges biological specimens, enabling the resolution of subdiffractional structures that would otherwise remain inaccessible^20^. Meanwhile, analytical approaches (e.g., SOFI^21^, SRRF^22^, MSSR^23^, DPR^24^) extract super-resolved information by statistically analyzing intrinsic fluorescent fluctuations^25,26^. More recently, learning-based approaches have emerged as a promising avenue in SRM, employing deep neural networks and curated training datasets to reduce post-processing complexity and computational burden^27,28^.

Despite these significant developments, SRM methodologies continue to encounter lingering challenges and unmet demands. For example, while deterministic techniques such as localization-based SRM can achieve remarkably nanometer-scale resolution, its extension into three dimensions often necessitates encoding axial information into single-molecule point spread functions (PSFs)^29-31^. However, such modifications are typically implemented using diffractive or adaptive optical elements, an approach that may compromise photon efficiency, provide marginal improvement, or increase instrumental complexity^32^. In contrast, deep learning-based methods enable streamlined computational workflows and enhanced accessibility across a broader range of experimental platforms^6,33^. Yet, these approaches frequently depend on substantial, high-quality training datasets that are often challenging to obtain or may not be readily accessible for novel biological discoveries^34^. For these reasons, further developments that simplify instrumentation, optimize photon utilization, and reduce data dependency, effectively integrating the strengths of both deterministic and learning-based paradigms, remain critically important for achieving more robust, versatile, and user-friendly SRM solutions.

In this study, we introduce deep volumetric localization microscopy (VLM), a super-resolution methodology for high-content single-molecule imaging through synergistic instrumental and algorithmic advances. Specifically, VLM captures single-molecule images with enhanced photon efficiency and precision through an optimally segmented optical aperture in conjunction with a Fourier light-field configuration. A cascaded neural network architecture then reconstructs the 3D sample geometry and extracts molecular coordinates with high accuracy. Crucially, VLM is trained exclusively on system-aware PSFs, eliminating dependencies on external imaging modalities or sample-specific training data. We validate VLM across diverse biological specimens, demonstrating its hardware simplicity, data-efficient operation, and low phototoxic impact. By synergizing deterministic and learning-based strategies, we anticipate VLM will empower fluorescence microscopy to overcome current imaging and computational limitations and drive broad advancements in biomedical research.

## RESULT

### The Principle and Framework of VLM

The VLM system leverages a high-content single-molecule imaging platform and an end-to-end neural network processing pipeline (**Fig. 1**). In practice, VLM acquires single-molecule images using a Fourier light-field microscope^35^, which effectively exploits aperture partitioning, depth extension, and computational microscopy^36-39^ (**Fig. 1a**). Specifically, an epi-fluorescence microscope is modified with a customized microlens array placed at the aperture plane, generating multiple perspective views tailored for single-molecule detection (**Methods**). Advancing previous prototypes^40-42^, the design in VLM improves photon budget, field of view, and spatial-frequency sampling, all critical considerations for optimum single-molecule data acquisition and processing (**Supplementary Figs. 1-3**).

**Figure 1.**
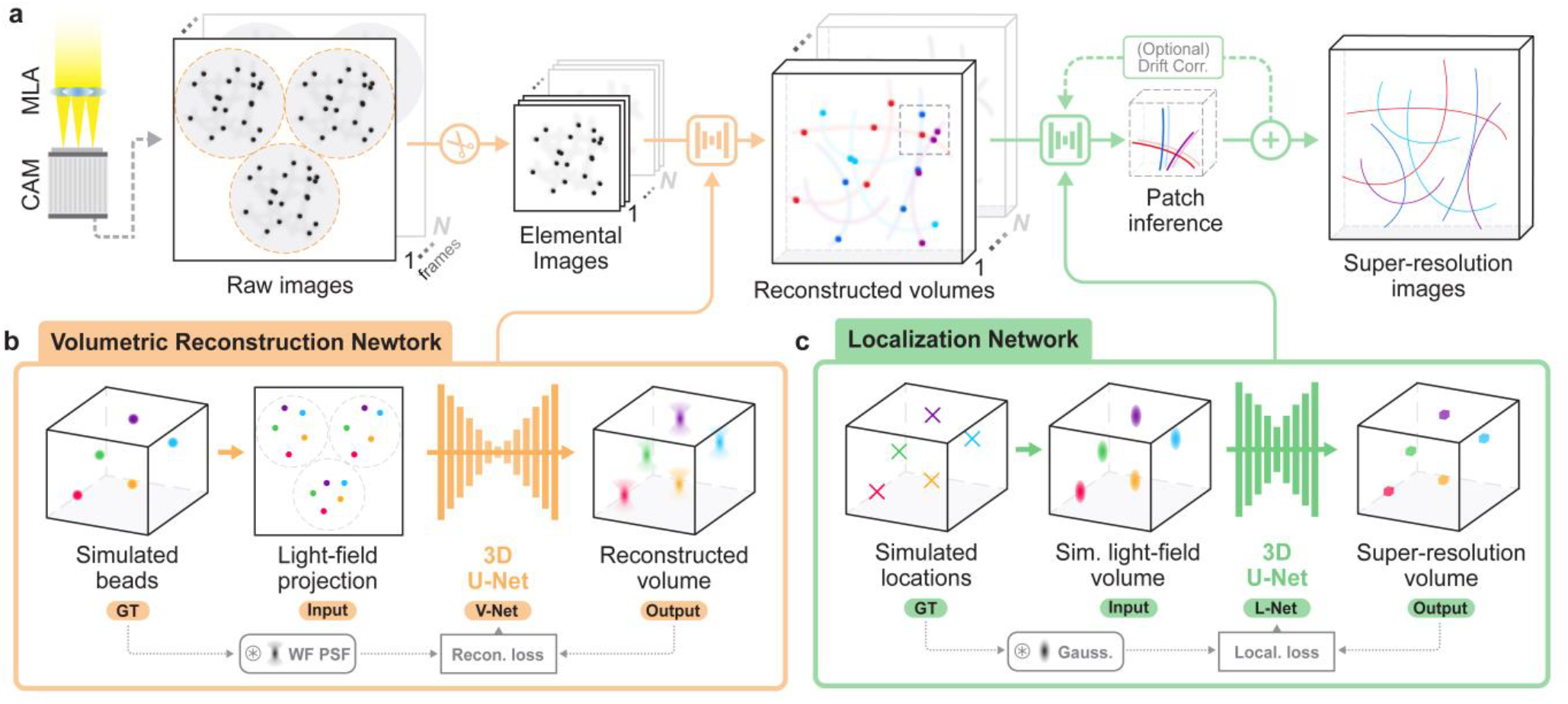
Deep volumetric localization microscopy (VLM). (a) Imaging and processing pipeline of VLM. Single-molecule image sequences are captured with a high-resolution Fourier light-field microscope. Raw frames are preprocessed, cropped, and fed into the volumetric reconstruction network (V-Net) for 3D reconstruction. Output volumes are partitioned into sub-regions for processing with the localization network (L-Net) and then concatenated into final super-resolved volumes. Optional drift correction ensures spatial fidelity. MLA, microlens array; CAM, camera sensor. (b) Schematic of V-Net, trained to decode 3D emitter positions from 2D light-field projections. Inputs (light-field projections of 3D-distributed simulated beads) are transformed into reconstructed 3D volumes, with reconstruction loss calculated against ground-truth wide-field point-spread functions (PSFs). (c) Schematic of L-Net, trained to predict super-resolved 3D emitter locations. Inputs are formed by convolving ground-truth simulated emitter positions with 3D Gaussian ellipsoids that match V-Net output resolution, with localization loss computed between predicted super-resolution and ground-truth volumes.

The recorded raw single-molecule sequences are processed into the tandem algorithmic pipeline, which synergistically integrates two convolutional 3D U-Net architectures (**Fig. 1, Methods, Supplementary Notes 1 and 2**). *First*, a volumetric reconstruction network (V-Net) recovers the 3D volume of fluorescent signals from 2D light-field data (**Fig. 1b**), a procedure conventionally performed using iterative methods (e.g., Richardson-Lucy deconvolution^43-45^). Unlike prior deep-learning approaches that require paired training data from high-end modalities (e.g., confocal or light-sheet microscopy)^46-51^, V-Net is trained exclusively on the native system-aware light-field PSF (**Supplementary Note 1 and Supplementary Table 1**). This strategy mitigates complex protocols, cross-modal alignment, and sample-specific constraints, streamlining workflows while ensuring broad applicability across biological specimens. *Next*, a localization network (L-Net) refines the 3D output volume of V-Net to pinpoint molecular positions and intensities (**Fig. 1c**). In contrast to previous methods that infer 3D positions reliant on 2D projections of engineered PSFs^52,53^, L-Net operates directly on volumetric data based on intrinsic 3D PSFs, achieving voxel-to-voxel super-resolution localization (**Supplementary Table 2**). This strategy accommodates refinement variants, thereby permitting an adaptive balance between computational efficiency and resolution demands (**Supplementary Note 2**). Critically, both networks rely solely on system responses (i.e., the PSFs), ensuring high data efficiency and usability without additional imaging modalities or sample-specific retraining processes.

### Characterization of VLM

To characterize VLM in biological samples, we first imaged immunostained β-tubulin in HeLa cells using 647-nm laser excitation and captured single-molecule images at 100 frames per second (fps) under continuous illumination (**Methods**). Compared with conventional wide-field microscopy, VLM acquired and processed single-molecule datasets across a FOV of 70 µm × 70 µm and an imaging depth of 5-10 µm, providing >10× higher resolution in all three dimensions and >5× extended imaging depth (**Fig. 2(a, b)**). The results resolved microtubule networks within densely packed regions, showing increased volumetric clarity and sectioning ability compared to standard Fourier light-field microscopy (**Fig. 2(c)**). Quantitative analysis of individual microtubule filaments revealed lateral and axial FWHM values of ∼60 nm and ∼90 nm, respectively (**Fig. 2(d)**). This quantification implied an effective 3D resolution near 20 nm laterally and 60 nm axially, considering the 50-60 nm width of immunostained microtubules^54^. These measurements correspond to localization precisions of approximately 10 nm laterally and 25 nm axially, consistent with those derived from single-molecule clusters (**Supplementary Fig. 4**).

**Figure 2.**
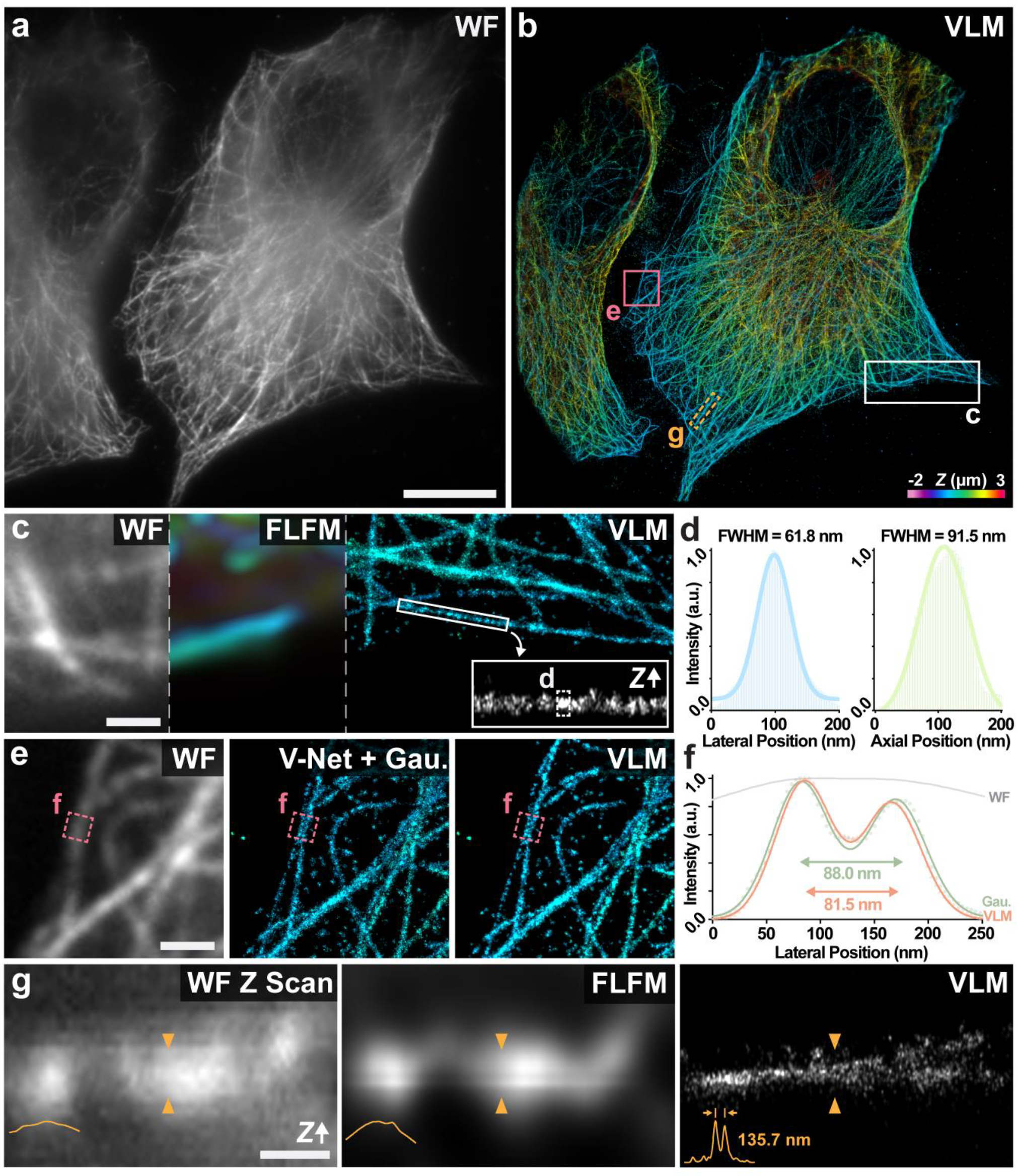
Imaging *β*-tubulin in HeLa cells using VLM. (a) Wide-field (a, *Z* = 0 µm) and 3D VLM (b) images of AF647-labeled *β*-tubulin. Z positions in (b) are color-coded according to the color-scale bar. Zoomed-in wide-field (left), Fourier light-field (middle), and VLM (right) image montage of the corresponding boxed region in (b). The inset shows the axial view of the boxed microtubule filament. Lateral (left) and axial (right) cross-sectional profiles of the filament as indicated in (c, inset), showing FWHM values of 61.8 nm and 91.5 nm, respectively. (e) Zoomed-in wide-field (left), V-Net + 3D Gaussian fitting (middle), and VLM (right) images of the corresponding boxed region in (b). (f) Lateral profiles of two microtubules indicated in (e) resolved by VLM and Gaussian fitting, showing a consistent 80-90 nm separation. (g) Wide-field (left), Fourier light-field (middle), and VLM (right) axial views of the corresponding boxed region in (b). Cross-sectional profiles show resolved microtubules 135.7 nm apart axially using VLM. Scale bars: 10 μm (a, b), 300 nm (c, e), 1 μm (g).

We further evaluated and compared the performance of L-Net with conventional single-molecule localization methods based on 3D Gaussian fitting^55^. Both approaches produced super-resolution volumes with high structural fidelity, resolving comparable non-specific substrate-binding molecular clusters (**Fig. 2e**). VLM distinguished microtubule filaments spaced 80-90 nm apart and at a distance of 130-140 nm, corresponding to an effective resolution of 20-30 nm and ∼60 nm, respectively, in the lateral and axial dimensions (**Fig. 2f, g**). The resolution measurements were validated using image decorrelation analysis^56^, which also confirmed that L-Net moderately outperformed 3D localization with Gaussian fitting, particularly across a ∼2 µm extended axial depth enabled by the volumetric reconstruction of V-Net (**Supplementary Fig. 5**). Compared to prior light-field-based single-molecule techniques^40^, VLM offers ∼4× improvement in lateral precision and nearly 2× improvement in axial precision, achieved using three elemental images at high acquisition speeds and at least 100× faster processing times using neural networks (**Supplementary Table 3**).

### Super-Resolution Imaging of Subcellular Components with VLM

Intracellular activities and dynamics are critical in regulating cellular functions, yet much of this information is obscured by the diffraction limit, making it inaccessible with conventional imaging techniques. Here, we demonstrated 3D super-resolution imaging of intracellular activities using VLM in fixed and live-cell samples. *First*, we imaged clathrin-coated pits (CCPs) in U-2 OS cells, which are essential for receptor-mediated endocytosis. During this process, clathrin proteins form a lattice-like coat to facilitate vesicle formation and cargo internalization^57^. In comparison to wide-field microscopy, VLM clearly visualized sub-diffraction-limited CCPs distributed within the 3D cellular environment (**Fig. 3a, b**). In particular, CCPs undergo various developmental stages, including nucleation, invagination, and scission^58^ (**Fig. 3c**). Using VLM, we captured these morphological changes and resolved hollow CCP structures measuring 50-120 nm in lateral diameter and 100-120 nm in axial diameter both at full maturity (**Fig. 3d**). *Next*, we utilized VLM to image MitoTracker diffused in live HeLa cells (**Fig. 3e**). MitoTracker is a photoswitchable membrane probe that enables single-molecule imaging of mitochondria without complex sample preparation^59^. Compared to conventional light-field deconvolution, VLM allows for resolving molecular locomotion surpassing the diffraction limit in all three dimensions (**Fig. 3f**). By assembling single-molecular localizations over temporal rolling windows, VLM reveals the dynamic patterns associated with mitochondrial movements across a 2-3 μm thick cellular space over time (**Fig. 3g**). *Finally*, we employed VLM for 3D single-particle tracking of fluorescently tagged proteins or vesicles in live cell^60^. Specifically, we performed 3D imaging and tracking of lysosomes and peroxisomes in live U-2 OS cells. Two-color images of each component were sequentially recorded by synchronizing camera exposures under stroboscopic illumination by two alternating laser lines^61^ (**Fig. 3h**). VLM processed the light-field acquisition and retrieved the dynamic movements of both organelles spanning a 5-μm axial range (**Fig. 3i, j**). With its high spatiotemporal resolution, VLM recorded the merging events of two lysosomes over a time period of seconds (**Fig. 3k, l**) and resolved two nearby peroxisomes that were otherwise unresolvable due to the diffraction limit (**Fig. 3m, n**). Quantifying individual diffusion coefficients of the two organelles showed the highly dynamic and locally confined features of lysosomes (mean = 7.09 × 10^−3^ μm^2^ s^-1^) and peroxisomes (mean = 0.65 × 10^−3^ μm^2^ s^-1^), respectively, in good agreement with time-lapse observation (**Fig. 3o**).

**Figure 3.**
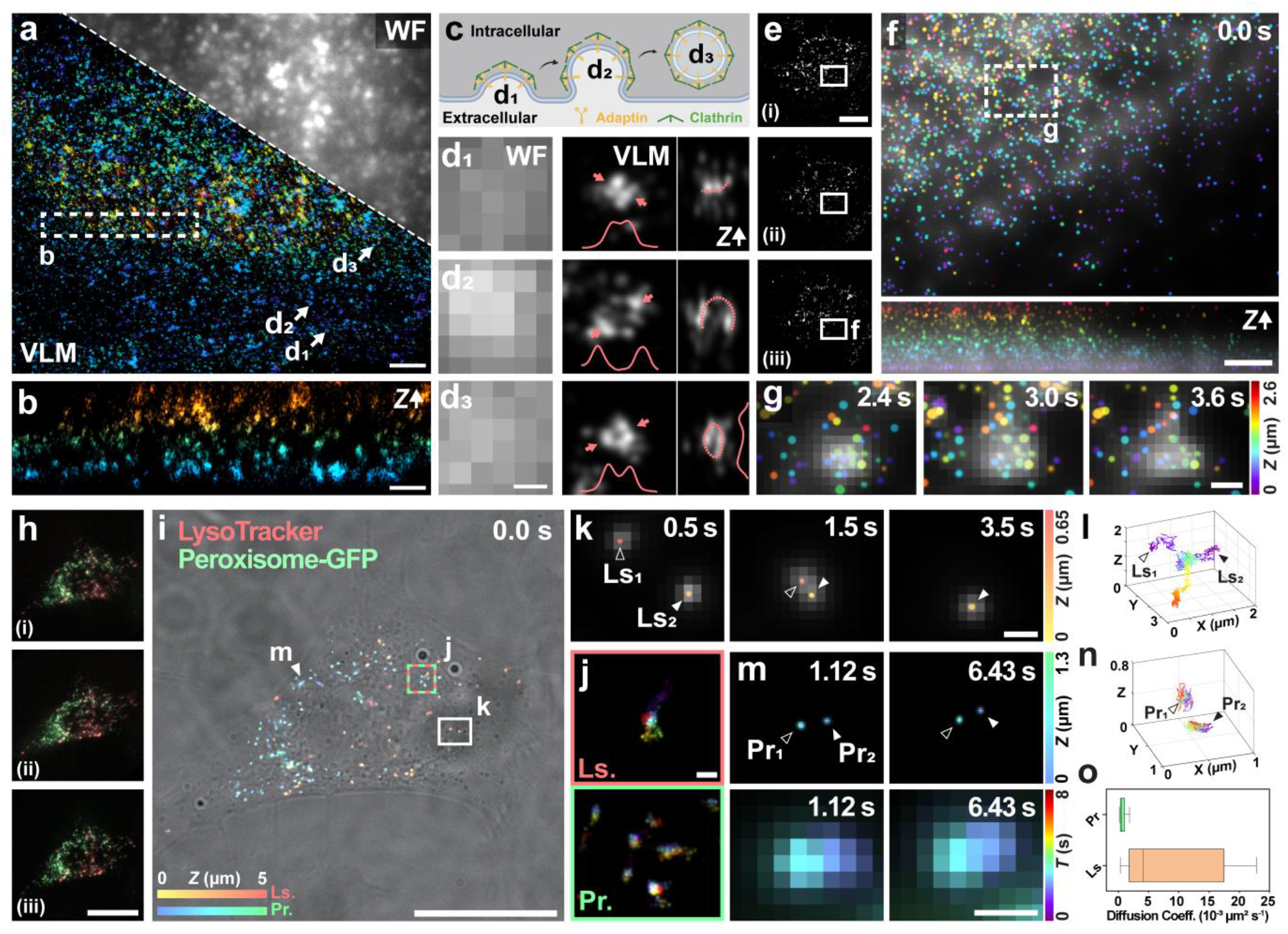
Volumetric imaging and dynamic tracking of intracellular processes using VLM. (a) Wide-field and 3D VLM images of AF647-labeled clathrin-coated pits (CCPs) in U-2 OS cells. (b) Cross-sectional projection of the corresponding boxed region in (a), revealing sparse 3D distribution of CCPs. (c) Schematic of CCP formation in three stages. (d) Wide-field (left) and 3D VLM (right) images of three representative CCPs, as indicated by arrows in (a), at distinct formation stages with cross-sectional profiles (insets). (e) Raw elemental light-field images of MitoTracker-labeled mitochondria in HeLa cells. (f) 3D VLM images of the boxed region in (e) at time point *t* = 0.0 sec, showing depth-color-coded emitters overlaid on corresponding light-field reconstructed images using Richardson-Lucy deconvolution. (g) Time-resolved MitoTracker dynamic changes through rolling-window analysis during high-power laser exposure at time points *t* = 2.4, 3.0, and 3.6 sec. (h) Raw elemental light-field images of lysosomes (red) and peroxisomes (green) in live HeLa cells. (i) Two-color 3D reconstructed image by V-Net (*t* = 0.0 sec) overlaid on the corresponding bright-field image. (j) Time-projected trajectories over 8 sec of lysosomes and peroxisomes in the boxed region in (i). (k) Zoomed-in images of two lysosomes (boxed in i), resolved by VLM at time points *t* = 0.5, 1.5, and 3.5 sec, overlaid on corresponding light-field reconstructed images. (l) 3D trajectories of the two lysosomes in (k) tracked over 8 sec. (m) VLM (top) and light-field reconstructed (bottom) images of two adjacent peroxisomes, as indicated in (i) at time points *t* = 1.12 and 6.43 sec. (n) 3D trajectories of the two peroxisomes in (m) tracked over 8 sec. (o) Comparative diffusion coefficients of lysosomes (mean = 7.09 × 10^−3^ μm^2^ s^-1^) and peroxisomes (mean = 0.65 × 10^−3^ μm^2^ s^-1^)). Scale bars: 2 μm (a, f), 100 nm (b, d), 20 μm (e, h, i), 500 nm (g, j-l).

### Super-Resolution Imaging of Staurosporine-Induced Cell Apoptosis

Apoptosis, the process of programmed cell death, is critical for maintaining normal biological function, while its dysregulation is frequently implicated in a variety of diseases^62^. The process is distinguished by hallmark morphological changes, including cell shrinkage, plasma membrane blebbing, and nuclear collapse^63,64^ (**Fig. 4a**). In addition, specific protein families have been identified as key regulators or indicators of apoptotic signaling. For instance, Bax, a pro-apoptotic member of the Bcl-2 family, translocates to and permeabilizes the outer mitochondrial membrane upon activation during apoptotic events^65^. Staurosporine (STS), a broad-spectrum protein kinase inhibitor, is commonly employed to induce apoptosis in various cell types^66-68^. Various studies have explored Bax-mitochondria interactions in apoptotic cells, primarily relying on conventional microscopy platforms^69-75^, which may overlook subtle molecular interactions occurring in 3D cellular contexts with finer details.

**Figure 4.**
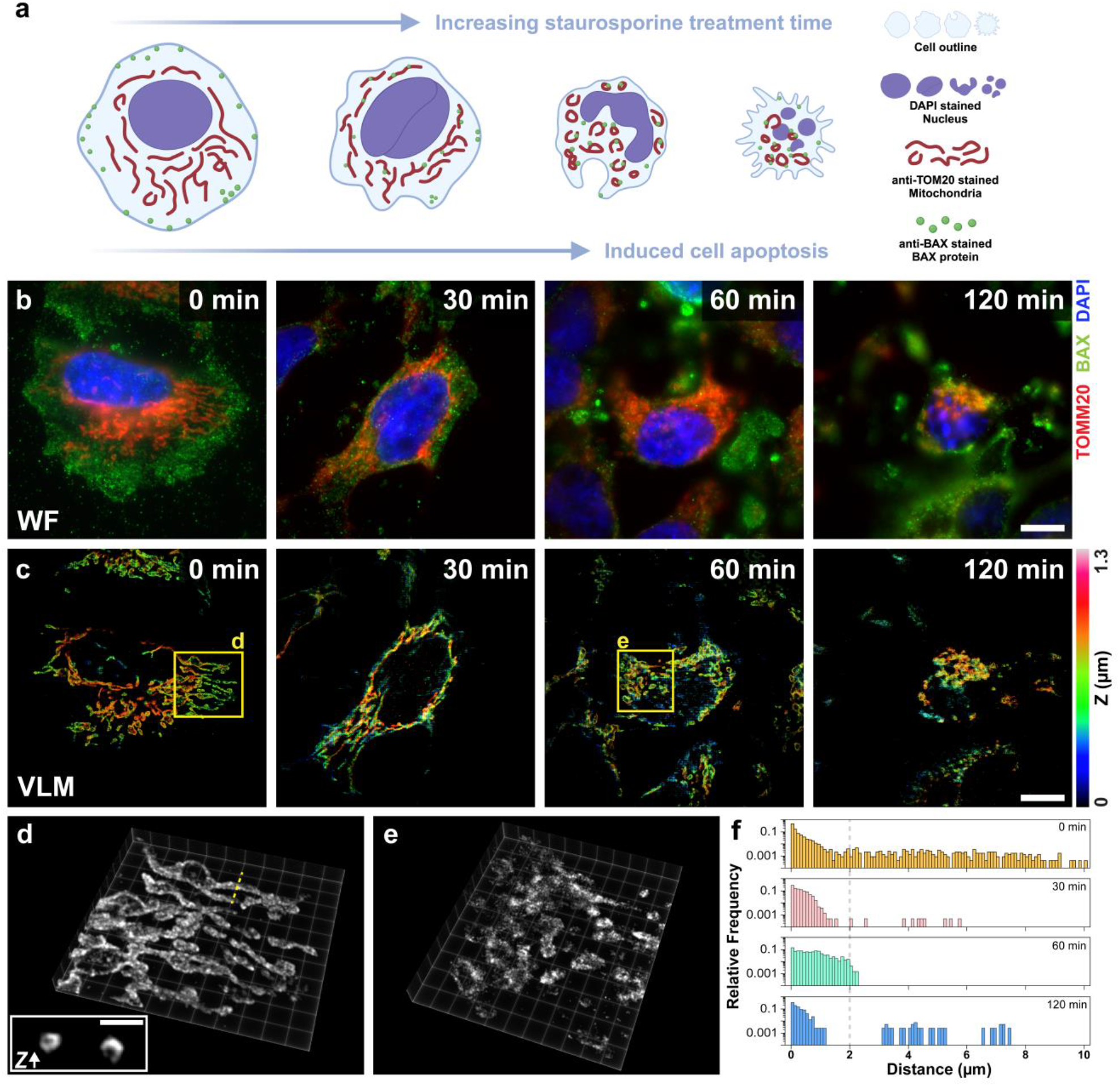
Super-resolution imaging of staurosporine-induced apoptosis in HeLa cells. (a) Schematic of apoptotic progression triggered by STS, showing key morphological changes, including cell shrinkage, membrane blebbing, nuclear collapse, and BAX protein translocation to mitochondria. (b) Wide-field images of cells treated with STS for 0, 30, 60, and 120 minutes, labeled with DAPI (nucleus, blue), AF488-BAX (green), and AF647-TOMM20 (mitochondria, red). (c) 3D VLM images of mitochondria corresponding to the treatment durations, revealing time-dependent structural remodeling. The depth information is coded according to the color scale bar. (d) Zoomed-in 3D view of mitochondria in (c, 0 min group) with axial cross-section (inset) showing hollow linear morphologies (diameters: left, 240 nm and right, 325 nm). (e) Circular mitochondrial structures in apoptotic cells (c, 60 min group), indicative of fragmentation during apoptosis. (f) Quantification of BAX-mitochondria proximity. The loss of long-distance BAX (>2 µm, dashed line) in treated cells confirms mitochondrial translocation. Scale bars: 10 μm (b, c), 1 μm (d, inset).

Here, we employed VLM to image 3D subcellular morphological alterations and Bax protein translocation in STS-induced apoptotic HeLa cells. In practice, HeLa cells were treated with 1-μM STS for 0, 30, 60, and 120 min, followed by wide-field and super-resolution imaging. Corresponding organellar and proteomic remodeling caused by STS treatment was observed in three-color wide-field imaging (**Fig. 4b, Supplementary Fig. 6**), revealing progressive nuclear condensation, mitochondrial fragmentation, and BAX translocation to mitochondrial membranes or membrane blebs during apoptosis. In addition, cells exhibited pronounced shrinkage and developed membrane blebs at advanced stages, aligning with established apoptotic hallmarks^76^. Meanwhile, VLM images resolved time-dependent mitochondrial swelling and eventual rupture as apoptosis progresses^77^. As observed, mitochondria without STS treatment exhibited tubular, hollow structures, whereas, after treatments of 60 min and 120 min, they displayed altered shapes and lost their distinctive features (**Fig. 4c-e**). Notably, VLM remained viable as the activation laser was restricted to ∼2 W/cm^2^, a decrease by at least an order of magnitude compared to typical STORM protocols^78,79^. This low-photon robustness of VLM not only minimized spectral crosstalk from the DAPI emission but also substantially reduced photodamage to the samples for long-term observation.

Finally, quantitative analysis of Bax-mitochondria proximity revealed dynamic redistribution during apoptosis (**Fig. 4f**). In the control group (0 min), most Bax localized >2 µm from mitochondria, indicating their initial association with the plasma membrane. As apoptosis advances, Bax proteins translocated toward mitochondria, evidenced by the disappearance of these long-distance events at 30 and 60 min. At later stages, some Bax proteins become sequestered in apoptotic bulges, reinstating longer distances to mitochondria within the contracted cell body. These observations underscore the capacity of VLM to reveal intricate subcellular morphological transformations and protein redistributions during apoptosis in 3D space^80^.

## DISCUSSION

Super-resolution microscopy, particularly localization-based methods, requires carefully balancing optical complexity, computational resources, and user accessibility^32^. Traditional approaches often prioritize either deterministic or learning-based methods but rarely unify their complementary strengths^18,81,82^. In this study, the VLM system significantly advances super-resolution microscopy by offering a streamlined platform for volumetric single-molecule imaging with an end-to-end neural network pipeline. This approach reduces instrumental complexity, facilitates network training under instrument-friendly conditions, and achieves efficient volumetric data reconstruction. These capabilities effectively enable the analyses of subcellular morphology, intracellular dynamics, and protein behaviors within the complex cellular microenvironment. The functionality of VLM can be further extended with novel fluorescent probes^83-86^, optical configurations^31,87-89^, and computational frameworks^90-94^. Furthermore, the common epi-fluorescence platform adopted by the VLM approach permits feasible integration to address broader biological discoveries with high-throughput systems^95,96^, single-molecule FRET^97^, and spatial-resolved transcriptomics^98,99^. We anticipate that VLM will serve as a powerful paradigm for elucidating the fundamental morphology and dynamics of complex biological systems beyond the optical and computational limit.

## METHODS

### Image Acquisition

The high-resolution Fourier light-field microscopy system (**Supplementary Fig. 1**) was developed using an epi-fluorescence microscope (Eclipse Ti2-U, Nikon Instruments), as narrated in the previous study ^35^. Briefly, an oil-immersion objective lens featuring 100× magnification and a numerical aperture (NA) of 1.45 (CFI Plan Apochromat Lambda 100× Oil, Nikon Instruments) was used. A piezo nano-positioner (Nano-F100S, Mad City Labs) was utilized for precise positioning. Samples were excited using multicolor laser lines (488 nm, 561 nm, 647 nm, MPB Communications), with the fluorescence collected through a quadband dichroic mirror (ZT405/488/561/647, Chroma) and a corresponding emission filter (ZET405/488/561/647 m, Chroma). The sample stage incorporated a micro-positioning system (MS2000, Applied Scientific Instrumentation) for accurate placement. The native image plane of the objective lens was Fourier-transformed using a Fourier lens (*f*_*FL*_= 275 mm, Edmund Optics). A customized microlens array (*f*_*MLA*_= 117 mm, RPC Photonics) was placed on the back focal plane of the Fourier lens (Supplementary Note 1). The elemental images formed by each microlens were captured using an sCMOS camera (ORCA-Flash 4.0 V3, Hamamatsu Photonics, pixel size *P*_*CAM*_ = 6.5 µm). During STORM imaging, the excitation was tuned to Hilo mode, guiding the laser beam toward the edge of the back focal plane^100^. Notably, the off-center HILO illumination not only increases the signal-to-noise ratio but also moves the background of the system’s internal reflection out of the field of view (**Supplementary Fig. 2**).

### Architecture of VLM

VLM is composed of two closely connected modules, a volumetric reconstruction network (V-Net) and a localization network (L-Net), for super-resolution visualization. Both networks adopt 3D U-Net architecture as the framework, and each has its own emphasis. The V-Net consists of seven layers and takes a pre-processed Fourier light-field image stack, which has been split based on the microlens positions and outputs a reconstructed volume (**Supplementary Note 1**). This network replaces the traditional heavy Richardson-Lucy algorithm and decodes the volume from the light-field images in one step, avoiding the lengthy iterative process. The reconstructed volume is patched, zero-padded, and up-sampled for the localization stage of VLM. The L-Net retains a three-layer network and takes pre-processed sub-volumes to output super-resolution volume with a grid size two-fold smaller than the effective camera pixel size (**Supplementary Note 2**). This network can be further expanded to a five-layer network to further shrink the output grid size down to 16.25 nm laterally and 32.5 nm axially. The three-layer or the five-layer version of L-Net can be used interchangeably based on the specific super-resolution task, or these two versions can be combined sequentially for drift correction (**Supplementary Fig. 3 and Supplementary Note 3**).

### Training of VLM

The volumetric reconstruction network and localization network are trained with simulated datasets without using additional imaging modalities, setting this work apart from the previously reported methods. Due to the difference in emphasis and nature of the V-Net and L-Net, distinct training strategies are applied. Specifically, for V-Net, sub-diffraction limit size beads are generated within a limited field-of-view that mimics the actual imaging setup (**Supplementary Note 1**). The beads are first convolved with a light-field PSF and projected to 2D light-field images as the input of the training process. The beads are then convolved with wide-field PSF in 3D as the ground truth pair for the corresponding 2D light-field image. The loss function is computed between output volume by network and wide-field volume by convolution. For L-Net, a set of locations in 3D continuous space is called first (**Supplementary Note 2**). At each location, a Gaussian ellipsoid is placed and pixelated to simulate the single emitters in the reconstructed 3D space. The loss function is computed by comparing the network-generated super-resolution volume with the pixelated locations. We trained the network with multiple rounds and various learning rates with ADAM optimizer^101^.

### Single Particle Tracking Analysis

Following VLM reconstruction of single-molecule data, peroxisomes, and lysosomes are further temporally grouped into trajectories using an ImageJ plug-in TrackMate^102-107^. Depending on the reconstruction quality and imaging settings, users choose the tracking parameters, including dark frame numbers, linking distance, and track length. The diffusion coefficient was then calculated from each trajectory using custom-made MATLAB code with msdanalyzer^108^, a customized MATLAB class.

### STORM imaging buffer

Imaging buffers for both fixed cell imaging and live cell imaging are prepared following the existing protocol^79,109^. Fixed cell STORM imaging buffers were used for imaging microtubules, clathrin-coated pits, and fixed mitochondria. When imaging with an 8-well glass bottom µ-Slide, each well was filled with 7 µL GLOX, 7 µL of 2-mercaptoethanol, and 690 µL buffer B. GLOX was prepared with 14 mg glucose oxidase, 50 µL catalase, and 200 µL buffer A, which consists of 10 mM Tris and 50 mM NaCl in PBS. Buffer B consists of 50 mM Tris, 50 mM NaCl, and 10% glucose in PBS. A live-cell STORM imaging buffer was used to image live mitochondria. The buffer was prepared with DMEM supplemented with 2% glucose, 6.7% of 1 M HEPES (pH 7.4), and an oxygen scavenging system that consists of 0.5 mg/mL glucose oxidase and 40 µg/mL catalase.

### Preparation and imaging of *β*-tubulin in fixed HeLa cells

Fixed microtubule staining was performed with HeLa cells. Cells were cultured in DMEM with 10% FBS and 1% Penicillin-Streptomycin (Pen-Strep) under 37 °C and 5% CO_2_ condition. Once reaching ∼80% confluency, cells were passaged and cultured in an µ-Slide with DMEM in each well. Immunostaining was performed following the STORM sample preparation protocol. Briefly, each well of the slide was first washed with PBS once. Then, each well was fixed with 3% PFA and 0.1% glutaraldehyde in PBS at room temperature for 12 minutes. Extra aldehyde groups were reduced by 0.1% sodium borohydride for 7 minutes, followed by 3 PBS washing, 5 minutes each. After that, cells were permeabilized and blocked with a blocking buffer (3% BSA with 0.2% Triton X-100 in PBS) for 30 minutes at room temperature. Cells were incubated for 30 minutes with β-tubulin primary antibody dilutions (BT7R, 10 µg/mL) in a blocking buffer at room temperature. Next, each well was washed 5 times with washing buffer (0.2% BSA with 0.05% Triton X-100 in PBS) for 15 minutes per wash at room temperature. After washing, labeled secondary antibody dilutions (goat anti-mouse, Alexa Fluor 647, 3 µg/mL) in blocking buffer were added to each well and incubated for 30 minutes at room temperature, with light avoided. Then, each well was washed 3 times with a washing buffer for 10 minutes per wash at room temperature, followed by one wash in PBS for 5 minutes. For better quality fluorescence imaging, cells were postfixed with 3% PFA and 0.1% glutaraldehyde at room temperature for 10 minutes, followed by 3 times washing in PBS for 5 minutes per wash. Finally, cells were stored in PBS for imaging purposes. Before STORM imaging started, the solution in the prepared wells was exchanged for fixed cell STORM imaging buffer, and the illumination of lasers was adjusted to HILO mode.

### Preparation and imaging of clathrin-coated pits (CCP) in fixed U-2 OS cells

Fixed CCP staining was performed with U-2 OS cells. The cells were cultured in DMEM with 10% FBS and 1% Penicillin-Streptomycin (Pen-Strep) under 37 °C and 5% CO_2_ conditions. Once reaching ∼80% confluency, cells were passaged and cultured in an 8-well glass-bottom µ-Slide with DMEM in each well. Immunostaining was performed using microtubule staining with brief modifications. Specifically, the permeabilization and blocking time was adjusted to 120 minutes at room temperature. The cells were incubated for 60 minutes with anti-clathrin primary antibody dilutions (EPR12235(B), 4 µg/mL) in a blocking buffer at room temperature. After washing, labeled secondary antibody dilutions (goat antirabbit, Alexa Fluor 647, 3 µg/mL) in blocking buffer were added to each well and incubated for 30 minutes at room temperature, light avoided. Then, each well was washed 3 times with a washing buffer for 10 minutes per wash at room temperature, followed by one wash in PBS for 5 minutes. For better quality fluorescence imaging, the cells were post-fixed with 4% PFA at room temperature for 10 minutes, followed by 3 times washing in PBS for 5 minutes per wash. Finally, cells were stored in PBS for imaging purposes. Before STORM imaging started, the solution in the prepared wells of µ-Slide was exchanged to fixed cell STORM imaging buffer, and the illumination of lasers was adjusted to HILO mode.

### Preparation and imaging of MitoTracker in live HeLa cells

Live mitochondria are stained and imaged with HeLa cells. The culturing protocol for HeLa cells is the same as described above. Once reaching the desired confluency, cells were passaged and cultured into a 35 mm imaging dish. The staining protocol followed the live cell imaging protocol with slight modification. Briefly, on the day of imaging, cells were incubated in 0.5 µM MitoTracker Deep Red in DMEM for 30 sec, washed 2 times with DMEM, and immediately used for imaging. During imaging, the objective heater was turned on, the illumination of lasers was adjusted to HILO mode, and the solution was exchanged for a live-cell imaging medium as described previously.

### Labeling of lysosome and peroxisome in live U-2 OS cells

Lysosome and peroxisome tracking was performed with live U-2 OS cells. The culturing protocol for HeLa cells is the same as described above. On the day prior to the imaging session, cells were passaged to a 35 mm imaging dish and cultured in DMEM with 20 µL of CellLight Peroxisome-GFP solution. Cells were then incubated under 37 °C and 5% CO_2_ condition for 18 hours. Next, cells were washed 2 times with DMEM and incubated with 100 nM LysoTracker Deep Red for 1 hour. Once the incubation was completed, the solution was exchanged for DMEM, and the dish was immediately used for imaging. While imaging, the objective heater was turned on, and the single particle tracking data was acquired in the epi-mode with two lasers (488 nm and 647 nm), alternatively illuminating the FOV.

### Induced apoptosis by staurosporine treatment of HeLa cells

HeLa cells were passaged and cultured in 8-well glass-bottom µ-Slide as described above. The slide was pre-treated with 0.1% Gelatin solution for 1 hour. On the imaging day, 1 μM of pre-warmed STS in DMEM was added to 3 wells, incubating 30, 60, and 120 min, respectively, at 37 °C. One additional well was reserved for non-treatment. The following procedures for preparing cells in the 4 wells are the same. After the treatment, cells were immunostained with TOMM20 (4 µg/mL) and Bax (4 µg/mL) primary antibody simultaneously for 30 minutes. TOMM20 was then labeled by an anti-rabbit secondary antibody conjugated with Alexa Fluor 647 (3 µg/mL), and Bax was labeled by an anti-mouse secondary antibody conjugated with Alexa Fluor 488 (3 µg/mL). The fixation, blocking, permeabilization, washing, and post-fixation steps were the same as the immunostaining protocol for microtubules. Finally, the slide was stained with DAPI dilutions (1 µg/mL) for 5 minutes. Each well was rinsed with additional PBS for storage. Before STORM imaging started, the solution in the prepared wells was exchanged for fixed cell STORM imaging buffer, and the illumination of lasers was adjusted to HILO mode.

### Resources, chemical, and biological materials

Figures 3c and 4a were created with BioRender.com. The sources of the chemicals and biological materials used in the experiments, including company names and catalog numbers, are listed in **Supplementary Table 4**.

## DATA AVAILABILITY

Data underlying the results presented in this paper may be obtained from the corresponding author upon reasonable request.

## CODE AVAILABILITY

The code is written in MATLAB (tested in 2023b, MathWorks) and Python 3.11. The latest version of the software is available at https://github.com/ShuJiaLab/VLM.

## ACKNOWLEDGEMENTS

We acknowledge the support of the National Institutes of Health grant R35GM124846 and the National Science Foundation grants EFMA1830941 and 2145235.

## AUTHOR CONTRIBUTIONS STATEMENT

K.H., X.H., and S.J. conceived and designed the project. K.H. and X.H. contributed to the construction of the optical system. K.H., X.H., and T.Q. conducted image processing and developed neural networks. K.H., Z.G., and X.W. prepared biological samples. K.H., T.Q., and X.W. performed imaging experiments. S.J. supervised the overall project. K.H. and S.J. wrote the manuscript with input from all authors.

